# Preexisting chronic infection skews the epigenetic landscape of subsequent memory CD8 T cell responses

**DOI:** 10.64898/2026.04.28.721334

**Authors:** Abdelhameed S. Dawood, Elizabeth E. Wolfson, Silmi Jiwani, Ahmet Ozdilek, Youssef M. Zohdy, Tylisha Gourdine, Sheikh Abdul Rahman, Jefferey A. Tomalka, Mohamed S. Abdel-Hakeem

## Abstract

Previous studies suggest that preexisting chronic disease impairs immune responses to subsequent infection and vaccination. However, the underlying epigenetic mechanisms are understudied. Here, we show that preexisting chronic infection with LCMV clone 13 (CL13) compromised the formation of central memory CD8 T cells (T_CM_) to subsequent *Listeria monocytogenes* infection, despite not profoundly impacting effector responses. This correlated with a skewed cytokine milieu. Our chromatin-accessibility profiling of Listeria-specific CD8 T cells showed significant epigenetic skewing of both T_CM_ and effector memory (T_EM_) in mice with preexisting LCMV-CL13, a skewing that started in memory-precursor effector cells (MPECs) during the effector phase. Transcription-factor binding sites (TFBS) analyses highlighted the interferon regulatory factor (IRF) family as major TFs implicated in this skewing. Thus, our results suggest that preexisting persistent inflammation skews the phenotypic and epigenetic landscape of subsequent memory responses, arguing for interventions to optimize vaccine-induced memory in hosts with preexisting chronic disease.

## Introduction

Vaccines are the most successful prophylactic measure against epidemics and pandemics, saving hundreds of millions of life and disability years globally over the past 100 years^1^. The main determinant of vaccine protective capacity is the formation of long-lived memory lymphocytes capable of mounting protective recall immune responses^2, 3^. Formation of neutralizing antibodies and generation of long-lived plasma cells and memory B cells (B_MEM_) are the main determinants of efficient and protective vaccine responses, respectively^4, 5, 6, 7, 8^. However, the generation of an efficient memory CD8 T cell (T_MEM_) response is another important correlate of protective immunity against intracellular pathogens, especially viruses like influenza and SARS-CoV-2^9, 10, 11^. This became more appreciated with emergence of SARS-CoV-2 Omicron variant, where mutations in the regions targeted by neutralizing antibodies (nAbs) (e.g. RBD in the spike region) rendered these nAbs inefficient in neutralizing this new strain. Only individuals with efficient memory CD8 T cell recall responses were protected from severe COVID-19, whereas individuals with less efficient T cell responses suffered from severe COVID-19 despite possessing high levels of nAbs^9, 10, 11^.

Numerous epidemiological studies suggest that people living with chronic diseases have increased susceptibility to acute infection, and might not be at the same level of protection upon vaccination (reviewed in^12^). Previous studies suggest that individuals living with chronic infections exhibit inferior immune responses to subsequent infections and vaccination^13, 14, 15, 16^ (and reviewed in^12, 17^). Chronic viral infection with HIV, HBV, HCV and HSV, or parasitic and helminths infections in endemic areas, e.g. plasmodium and Leishmania, were all associated with increased susceptibility and severity of unrelated infections. It has also become appreciated that patients with solid and hematologic cancer are more vulnerable to SARS-CoV-2 infection and have poorer prognoses of COVID-19 (reviewed in^18^), as well as lower responses to COVID-19 vaccine^19^. Similarly, people living with HIV (PLWH) show lower seroconversion in response to COVID-19 vaccine^20^. Recently, Faleti et al. have shown that individuals with SLE also had a compromised response to COVID-19 vaccine^21^. However, the underlying mechanisms of this compromised response to subsequent infections and vaccination are not completely understood.

A few studies in animal models aimed at examining the potential causes for inferior recall capacity by memory T cells in hosts with preexisting chronic infection. Examining the cellular profiles of antigen-specific T cells, confirmed the inferior expansion of T_MEM_ generated to acute infection in response to proinflammatory cytokine signals and in hosts with preexisting chronic infection (hereafter termed inflm-T_MEM_)^22, 23^. Transcriptional profiling of inflm-T_MEM_ in one study revealed enrichment of signatures associated with proinflammatory pathways, e.g. type I interferon (IFN-I), and TLR4/TLR7 signaling associated with TNF and IL-6^22^. The skewed phenotype and transcriptional signature were associated with compromised recall response and protective capacity, and the phenotype was partially restored by knocking-out IFN-I receptor (IFNAR) on antigen-specific CD8 T cells^22^. Transcriptional profiling of CMV-specific CD8 T_MEM_ from individuals living with chronic hepatitis C virus (HCV) infection displayed similar skewing of their transcriptional profiles compared to individuals with no known preexisting chronic diseases. Importantly, CMV-specific T_MEM_ from HCV+ humans displayed the same enriched signature for inflm-T_MEM_ from mice with preexisting chronic lymphocytic choriomeningitis virus (LCMV) with the clone 13 strain (LCMV-CL13)^22^. Few studies from other groups suggest that the general impact on T_MEM_ is similar, e.g. in the chronic lymphocytic leukemia (CLL) model^24^, and in preexisting helminths infections^25^. Martens et al. focused on the impact of preexisting chronic lymphocytic leukemia (CLL) on early time points at the peak of the effector OT-I response to mouse CMV expressing OVA (mCMV-OVA)^24^. Their data confirmed a skewing of the transcriptional and epigenetic profiles of antigen-specific CD8 T cells to acute infection in hosts with preexisting chronic disease, but these data were generated on total CD8 T cells that already had different subsets’ frequencies^24^.

However, the impact of preexisting chronic disease on the epigenetic programs of T_MEM_ subsets has not been studied. Here, we examined the impact of preexisting chronic viral infection on the programing of memory T cell subsets responding to subsequent acute infections. For this, we tracked the differentiation of pathogen-specific OT-I CD8 T cells in response to *Listeria monocytogenes* expressing ovalbumin (LM-OVA) in mice with preexisting chronic LCMV-CL13 infection compared to control mice with no preexisting infection. We confirmed and extended previous findings that preexisting chronic infection skewed the phenotype and subset dynamics of LM-OVA-specific OT-I cells during the early effector phase of the response, and through the memory differentiation stages, where the establishment of the central memory subset was specifically and significantly compromised. Examining the cytokine milieu in the serum at baseline, before Listeria infection, demonstrated significantly higher levels of several proinflammatory cytokines, including gamma-interferon (IFNΨ), IFNΨ-induced proteins 10 (IP-10/CXCL10), and tumor necrosis factor alpha (TNFα). Our chromatin-accessibility profiling of both the central memory (T_CM_) and effector memory (T_EM_) revealed significant skewing of the epigenetic programs of both T_CM_ and T_EM_ in mice with preexisting chronic infection. This epigenetic skewing originated early in the memory precursor effector cells (MPECs) subset, and many of its features become fixed in T_CM_, T_EM_, or both. Transcription-factor binding sites (TFBS) analyses of our ATACseq data highlighted the interferon regulatory factor (IRF) family as significantly impacted TFs. Together, our findings suggest that preexisting chronic diseases with persistent inflammatory cytokine milieu induces skewing of the cellular and subset profile of memory T cells to subsequent acute infection, as well as skewing of their epigenetic programs. These studies highlight the need for therapeutic approaches to modulate this skewing upon vaccinating individuals with preexisting chronic diseases, especially diseases with underlying persistent inflammation.

## Results

### Preexisting chronic infection does not inhibit effector responses to subsequent acute infection

Previous studies suggest that effector CD8 T cell responses to acute infection in mice with preexisting chronic disease, e.g. chronic lymphocytic leukemia (CLL), were skewed according to different metrics, including skewed subset dynamics of effector T cells (T_EFF_) and reduced production of functional cytokines, e.g. gamma-interferon (IFNγ), tumor necrosis factor (TNF), and expression of the degranulation marker LAMP-1/CD107a^24^.

We used a similar experimental design to previous studies^22, 24^, where C57/BL6 (B6) mice were infected with the chronic strain of LCMV, LCMV Clone 13 (LMCV-CL13), and after entering the chronic phase, 4 weeks post CL13-infection, the mice were subsequently infected with of *Listeria monocytogenes* expressing ovalbumin antigen (LM-OVA), 10^4^ CFU/mouse. One day prior to LM-OVA infection, 2×10^4^ congenically distinct OVA-specific OT-I CD8 T cells were adoptively transferred to the CL13-infected mice. As a control, a matched group of co-housed mice with no preexisting infection received the same number of OT-I cells and were infected with LM-OVA (Fig.1a). OT-I cells isolated from spleen were profiled at the peak of the effector immune response to LM-OVA, ∼8 days (7-9 days) post-infection with LM-OVA (hereafter termed d8).

**Figure 1:**
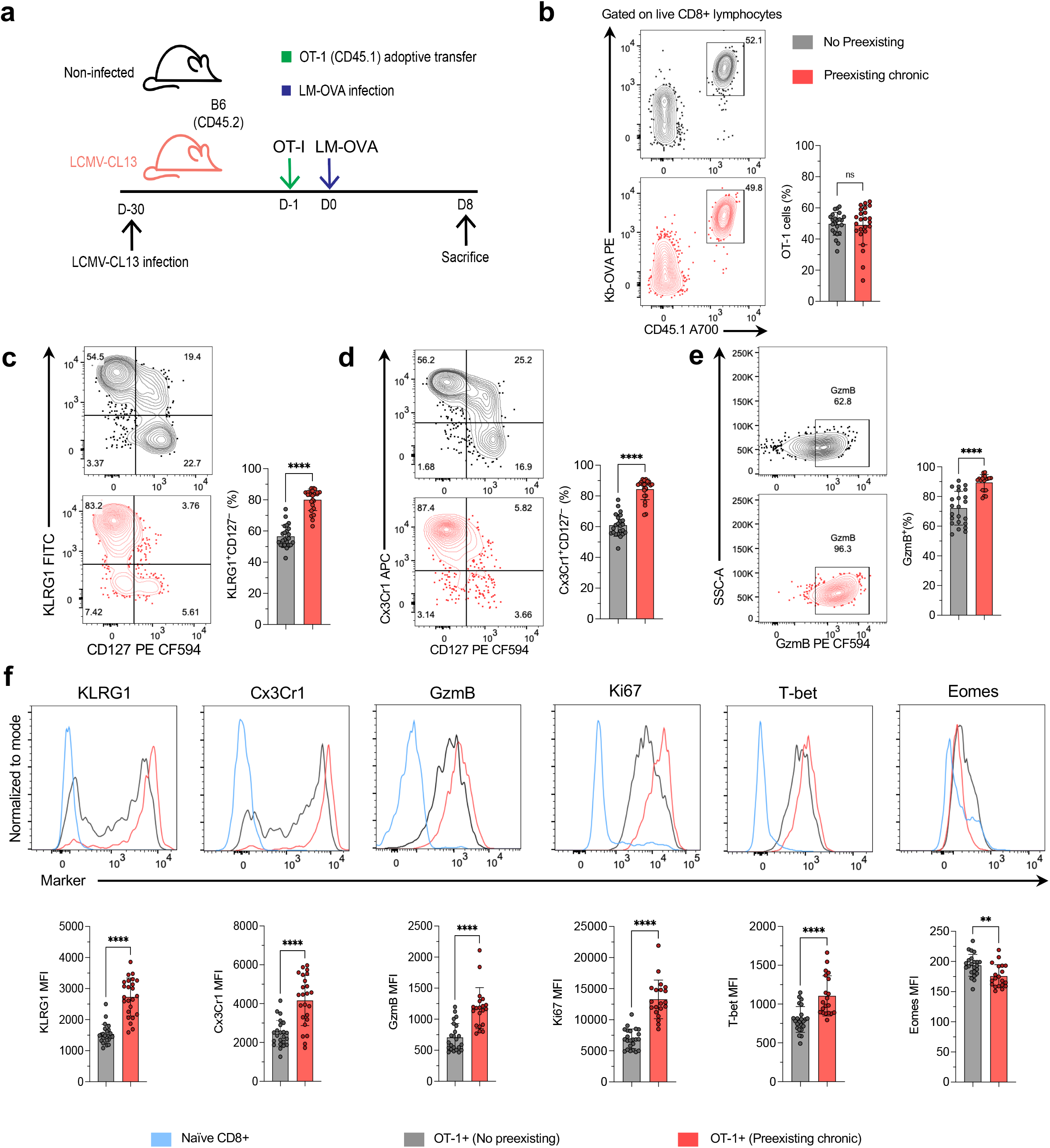
Preexisting chronic infection does not inhibit expansion or activation of effector T cells (Teff) to subsequent acute infection. (A) Experimental design. Mice without or with preexisting LCMV-CL13 chronic infection (top/grey and bottom/red, respectively) received 2×10^4^ congenic K^b^-OVA-specific transgenic CD8 T cells (OT-I) then infected with *Listeria monocytogenes* expressing ovalbumin (LM-OVA) and OT-I responses profiled 8 days (d8) post-infection with LM-OVA. (B) Left, representative dot plots showing OT-I frequency in splenocytes isolated from mice with preexisting chronic infection (bottom/red) compared to control mice without preexisting chronic infection (top/grey). Right, summary bar-plots. Gated on live CD8 T cells. (C-E) Left, representative dot plots showing frequency of LM-OVA specific OT-I cells expressing the different combination of markers. Right, summary bar-plots. (F) Top, representative histograms for level of expression of effector-associated markers on OT-I cells compared to naïve CD8 T cells (blue), represented as the mean fluorescence intensity (MFI). Bottom, summary bar-plots. (C-F) Gated on OT-I live CD8 T cells, or live CD8 T cells for naïve staining controls. Results from at least 3 independent experiments 2-5 mice/group. ** P<0.01, **** P<0.0001

Our data indicate that there was no significant difference in the frequency of LM-OVA-specific OT-I cells (OT-Is) at d8 (Fig.1b), unlike previous results that showed increased frequency and number of OT-Is responding to mouse cytomegalovirus (mCMV) in mice with preexisting CLL compared to non-CLL mice^24^. Our data showed significantly lower number of OT-I cells in the group with preexisting CL13-infection, but this was confounded by the fact that the total number of CD8 T cells and splenocytes were significantly lower in number in this group (Data not shown. Supplementary figures pending). However, similar to previous data^24^, we observed significant skewing of the main subsets of T_EFF_, the KLRG1+ CD127– short-lived effector cells (SLECs) and the CD127+ KLRG1– memory-precursor effector cells (MPECs), where SLECs were significantly enriched in mice with preexisting-CL13 (Fig.1c), together with higher expression levels of the effector associated transcription factor (TF) T-bet and lower expression of the memory-associated TF eomesdermin (Eomes) (Fig.1f).

We also extended previous findings by showing that the frequency and levels of expression of another marker of activation, Cx3cr1, and the effector molecule, granzyme B (GzmB) were significantly higher in mice with preexisting-CL13 (Fig.1d-e).

To examine simultaneous production of cytokines by OT-I cells, i.e. polyfunctionality, we evaluated the production of the effector cytokines IFNg, TNF, and IL-2, as well as expression of the degranulation marker CD107a/LAMP-1 upon ex-vivo re-stimulation with OVA peptide. Our data indicate significantly lower frequency of polyfunctional cells producing all four measured functions, cytokines IFNg, TNF, and IL-2, as well as CD107a expression in mice with preexisting-CL13 (Data not shown. Supplementary figures pending). However, this was balanced by higher frequency of triple-functional CD8 T cells in mice with preexisting-CL13, thus the total frequency of cells with 3-4 functions is not significantly different.

Previous studies showed that preexisting chronic CLL did not significantly impact the viral load of subsequent acute infection with mCMV evaluated on day 8 post mCMV-infection (d8)^24^, but whether the skewed responses would impact the kinetics of acute pathogen control was not thoroughly investigated. Examining the bacterial loads on d8 post-infection with LM-OVA low infection dose (1×10^4^ CFU/mouse) that was used throughout this study, showed clearance of Listeria from both the liver and spleen in all mice (data not shown). When examining the bacterial loads at earlier time points (d4) upon infection with 5-fold higher infection dose (5×10^4^ CFU/mouse), mice with preexisting-CL13 showed slightly lower bacterial loads in both the liver and spleen compared to mice with no preexisting infection (Data not shown. Supplementary figures pending).

Together, our results examining the profile of antigen-specific OT-I CD8 T cells to acute infection with LM-OVA in mice with preexisting LCMV-CL13 chronic infection confirmed previous results using mCMV acute infection in mice with preexisting CLL^24^ that showed increased frequency of SLECs on the expense of MPECs. We extended these previous findings by demonstrating that other markers of antigen-specific CD8 T cells were also significantly impacted.

### Preexisting chronic infection skews the memory responses to subsequent acute infection

Previous studies suggest that memory CD8 T cell responses to acute infection in mice with preexisting chronic infection, e.g. LCMV-CL13 or Toxoplasma, were skewed. Where the subset dynamics of memory T cells (T_MEM_) were skewed and reduced production of functional cytokines, e.g. gamma-interferon (IFN γ), tumor necrosis factor (TNF), and expression of the degranulation marker LAMP-1/CD107a^22^.

We confirmed and expanded these findings, where longitudinal analysis of LM-OVA-specific OT-I cells at the peak effector response (d8) were expanded to the memory phase, 4 weeks post Listeria infection (Fig.2a). Using the same experimental design we used for studying the effector phase, we examined the phenotype of Listeria-specific OT-I CD8 T cells at d30 post-Listeria infection.

**Figure 2:**
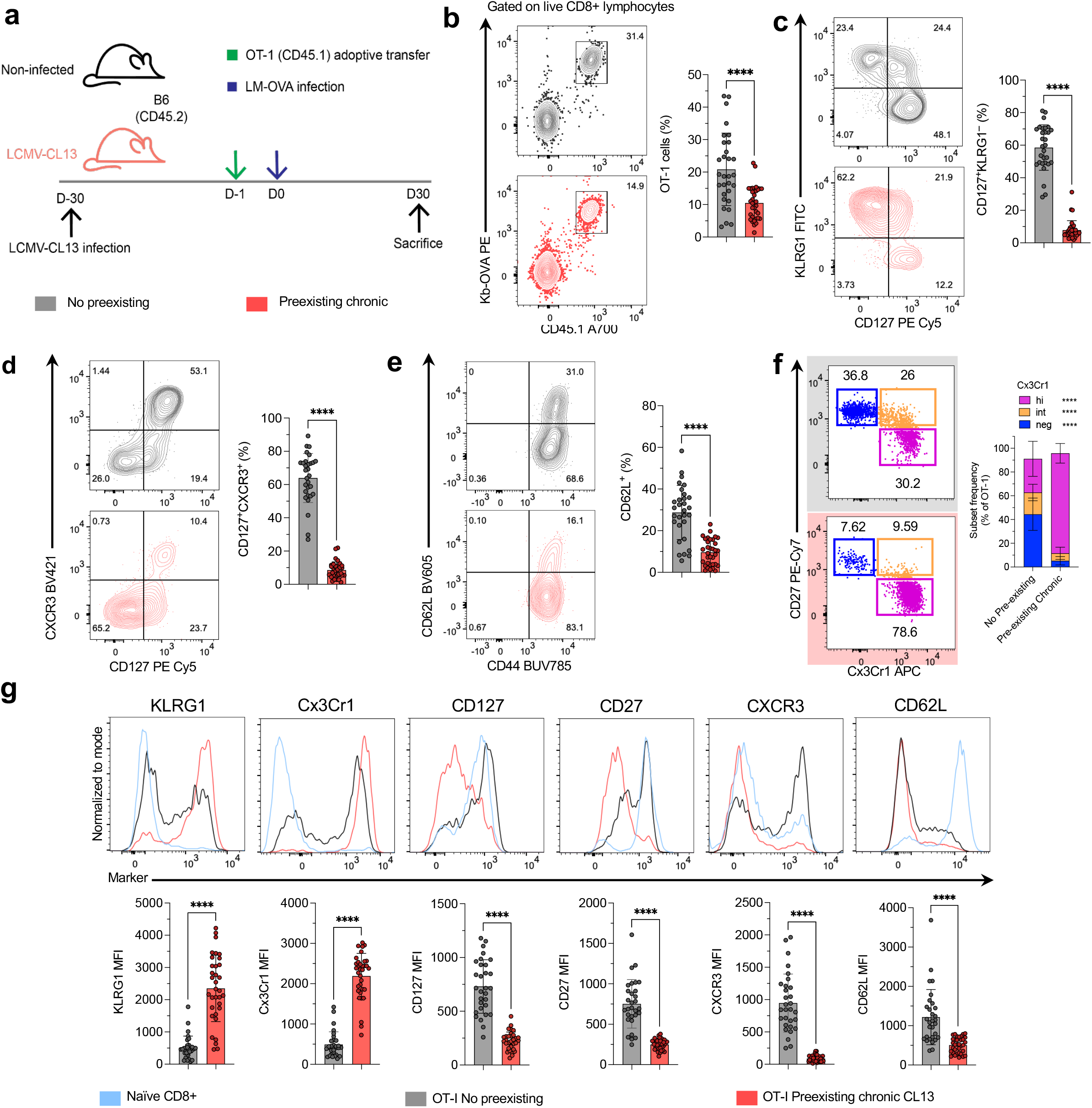
Preexisting chronic infection skews phenotype of memory T cells (Tmem) to subsequent acute infection. (A) Experimental design. Mice without or with preexisting LCMV-CL13 chronic infection (top/grey and bottom/red, respectively) received 5-20×10^3^ congenic K^b^-OVA-specific transgenic CD8 T cells (OT-I) then infected with *Listeria monocytogenes* expressing ovalbumin (LM-OVA) and OT-I responses profiled 30 days (d30) post-infection with LM-OVA. (B) Left, representative dot plots showing OT-I frequency in splenocytes isolated from mice with (bottom/red) or without preexisting chronic infection (top/grey). Right, summary bar-plots. Gated on live CD8 T cells. (C-F) Left, representative dot plots showing frequency of LM-OVA specific OT-I cells expressing the different combination of markers. Right, summary bar-plots. (G) Top, representative histograms for level of expression (MFI) of effector and memory-associated markers on OT-I cells compared to naïve CD8 T cells (blue). Bottom, summary bar-plots. (C-G) Gated on OT-I live CD8 T cells, or live CD8 T cells for naïve staining controls. Results from at least 3 independent experiments 2-5 mice/group. **** P<0.0001

Our results showed significant difference in the frequency of LM-OVA-specific OT-I cells (OT-Is) at d30 post infection, where the frequency of OT-I T_MEM_ was ∼2-fold lower in mice with preexisting-CL13 (Fig.2b). More significant was the skewing of LM-OVA-specific OT-I’s phenotype, where the frequency of CD127+ KLRG1– memory cells was ∼4-6-fold lower in mice with preexisting-CL13, with a mean <10%, compared to ∼60% in mice with no preexisting infection (Fig.2c). Overall, the frequency of total CD127+ OT-I cells was consistently lower (∼2-fold) in mice with preexisting-CL13 (Fig.2c and 2d).

Within T_MEM_ OT-I cells, mice with preexisting chronic LCMV-CL13 also showed significantly lower frequencies of cells expressing the memory marker CXCR3 (Fig.2d). Similarly, OT-I cells expressing the central memory marker CD62L cells were significantly lower in frequency in mice with preexisting LCMV-CL13 (Fig.2e). Using the T_MEM_ sub-setting by Gerlach et al.^26^, confirmed a significant decrease of the frequency of CD27+ CX3CR1– central memory (T_CM_) and the transitional memory CD27+ CX3CR1-intermediate, whereas the CD27– CX3CR1-hi effector memory (T_EM_) were significantly enriched in the group with preexisting chronic infection (Fig.2f).

The significant differences were not limited to the skewing of subset dynamics and frequency of different populations, but was also observed for the levels of expression of numerous T cell differentiation markers, where the mean fluorescence intensity (MFI) for OT-I expression of memory, central memory, and differentiation associated markers CD127, CXCR3, CD62L and CD27 was significantly lower for mice with preexisting chronic LCMV (Fig.2g)

Similar to our polyfunctionality data for d8, polyfunctionality data for d30 indicate that despite the lower frequency of polyfunctional cells producing all four measured functions cells in mice with preexisting-CL13, this group exhibited higher frequency of triple-functional CD8 T cells, leading to a total frequency of cells with 3-4 functions that is not significantly different (Data not shown. Supplementary figures pending).

Thus, using different markers for T cell memory and memory subsets, our data indicate that preexisting chronic infection significantly impacted the memory differentiation of antigen-specific CD8 T cells responding to subsequent acute infection. The major impact was observed for the central memory subset, although the phenotype and expression of memory-associated markers was impacted for total T_MEM_ OT-I cells.

### Central memory T cell formation remains compromised long-term post Listeria clearance in hosts with preexisting chronic infection persist

We next sought to confirm whether the skewing in phenotype and subset dynamics of T_MEM_ was temporary and would become less significant at later time points post clearance of the acute infection. For this, we tracked the LM-OVA-specific OT-I cells after prolonged durations post clearance of LM-OVA at d68-d100 post infection (d60+) (Fig.3a).

**Figure 3:**
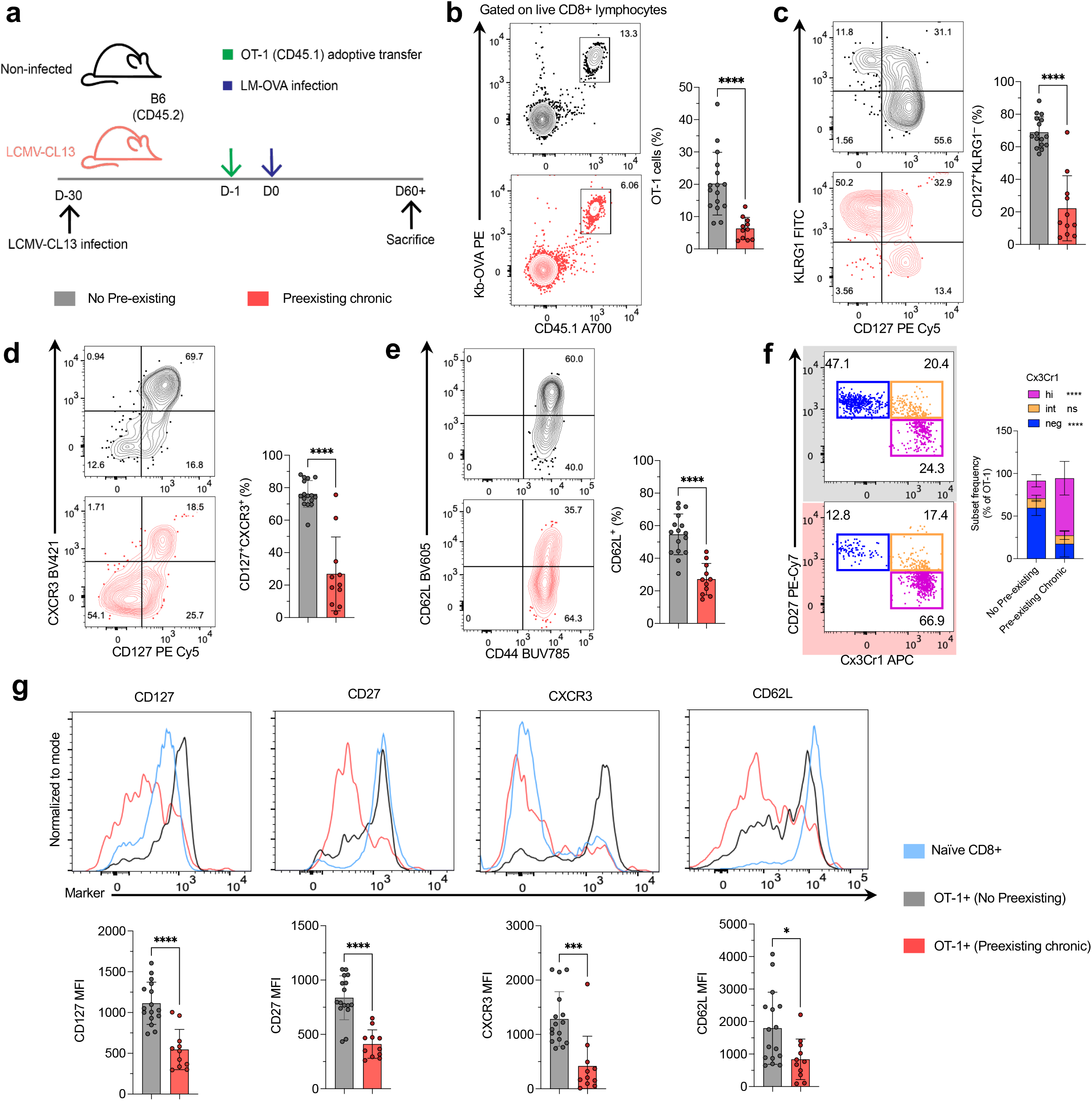
Skewed Tmem phenotype due to preexisting chronic infection persists long term. (A) Experimental design. Mice without or with preexisting LCMV-CL13 chronic infection (top/grey and bottom/red, respectively) received 5-20×10^3^ congenic K^b^-OVA-specific transgenic CD8 T cells (OT-I) then infected with *Listeria monocytogenes* expressing ovalbumin (LM-OVA) and OT-I responses profiled 68-100 days (d60+) post-infection with LM-OVA. (B) Left, representative dot plots showing OT-I frequency in splenocytes isolated from mice with (bottom/red) or without preexisting chronic infection (top/grey). Right, summary bar-plots. Gated on live CD8 T cells. (C-F) Left, representative dot plots showing frequency of LM-OVA specific OT-I cells expressing the different combination of markers. Right, summary bar-plots. (G) Top, representative histograms for level of expression (MFI) of effector and memory-associated markers on OT-I cells compared to naïve CD8 T cells (blue). Bottom, summary bar-plots. (C-G) Gated on OT-I live CD8 T cells, or live CD8 T cells for naïve staining controls. Results from at least 3 independent experiments 2-5 mice/group. * P<0.1, *** P<0.001, **** P<0.0001

Like the significant difference in frequency of OT-I cells at d30 post infection, the frequency of OT-I Tmem at d60+ was lower in mice with preexisting-CL13 (Fig.3b). In fact, the difference was even more significant than at d30 where the mean OT-I% was ∼4-fold lower in the group with preexisting chronic infection. Similarly, a significant percentage of OT-I cells remained in a KLRG1+ CD127– effector-like state in mice with preexisting chronic disease, whereas more OT-I cells from the control mice were in the memory CD127+ KLRG1– state (Fig.3c). Mice with preexisting chronic LCMV-CL13 still showed significantly lower frequencies of CXCR3+ cells (Fig.3d), CD62L+ cells (Fig.3e), and CD27+ CX3CR1– T_CM_ (Fig.3f).

Notably, more OT-I cells were differentiating into CXCR3+ CD127+, CD62L+, and CD27+ CX3CR1– T_CM_ at the time points d60+ compared to d30 in the control group, whereas the accumulation of higher frequencies of OT-I cells with a T_CM_ phenotype seemed compromised.

The levels of expression of memory and central memory associated markers CD127, CXCR3, CD62L and CD27 remained significantly lower for mice with preexisting chronic LCMV (Fig.3g)

Thus, our results indicate that T_MEM_ have an arrested phenotype, where the formation of T_CM_ was significantly compromised even after more than 60 days post Listeria infection.

### Skewed serum cytokine milieu in mice with preexisting chronic LCMV correlates with compromised central memory formation

Previous work has indicated a transcriptional profile of inflm-T_MEM_ that is enriched in proinflammatory signature. However, none of these studies examined the levels of cytokines in hosts with preexisting chronic disease. Thus, we next sought to investigate the cytokine milieu in the serum of mice with preexisting chronic LCMV-CL13 infection compared to mice with no preexisting infection.

Using the Mesoscale Discovery (MSD) platform, we measured the level of 19 cytokines: interferon alpha (IFN-α), IFN-β, IFN-γ, interleukin (IL)-1β, IL-2, IL-4, IL-6, IL-10, IL-12p70, IL-13, IL-17A/F, IL-21,IL-23, IFNγ-induced protein (IP)-10, keratinocyte chemoattractant/growth-regulated oncogene (KC/GRO), monocyte chemoattractant protein (MCP)-1, macrophage inflammatory protein-1 alpha (MIP-1α), MIP-1β, and tumor necrosis factor alpha (TNFα). Principal component analysis (PCA) of the levels of measured cytokines in the serum from mice with preexisting chronic LCMV-CL13 compared to control mice with no preexisting infection at baseline timepoint (1-3 days before *Listeria* infection) showed distinct profiles between the two groups (Fig.4a). This was consistently evident for mice from 3 different experiments for specific cytokines that seemed to drive this distinct cytokine profile from mice with preexisting CL13 versus control mice; MCP-1, IL-10, MIP-1α, and TNFα, followed by IFN-γ and KC/GRO, and to a lesser extent IL-6 and MIP-1β (Fig.4b-d). More importantly, when evaluating the correlation of the significantly different cytokine levels at baseline with the skewed phenotype observed for T_MEM_ OT-I on d30, there was a strong correlation between reduced frequencies of OT-I cells with a CD127+ CXCR3+ memory phenotype and the increased levels of cytokines, e.g. IFN-γ and TNFα (Fig.4e). Generally, this was true for several other cytokines, including IL-6, IL-10, and IP-10 (Fig.4f). Additionally, this skewed cytokine milieu was associated with decreased levels of expression of the memory markers CD127 and CXCR3, and the differentiation markers CD27 and CD44 (Fig.4f). Expectedly, the higher levels of these cytokines in mice with preexisting CL13 were also associated with increased frequency of KLRG1+ cells and higher levels of expression of effector-associated markers, KLRG1 and Cx3cr1 (Fig.4f).

**Figure 4:**
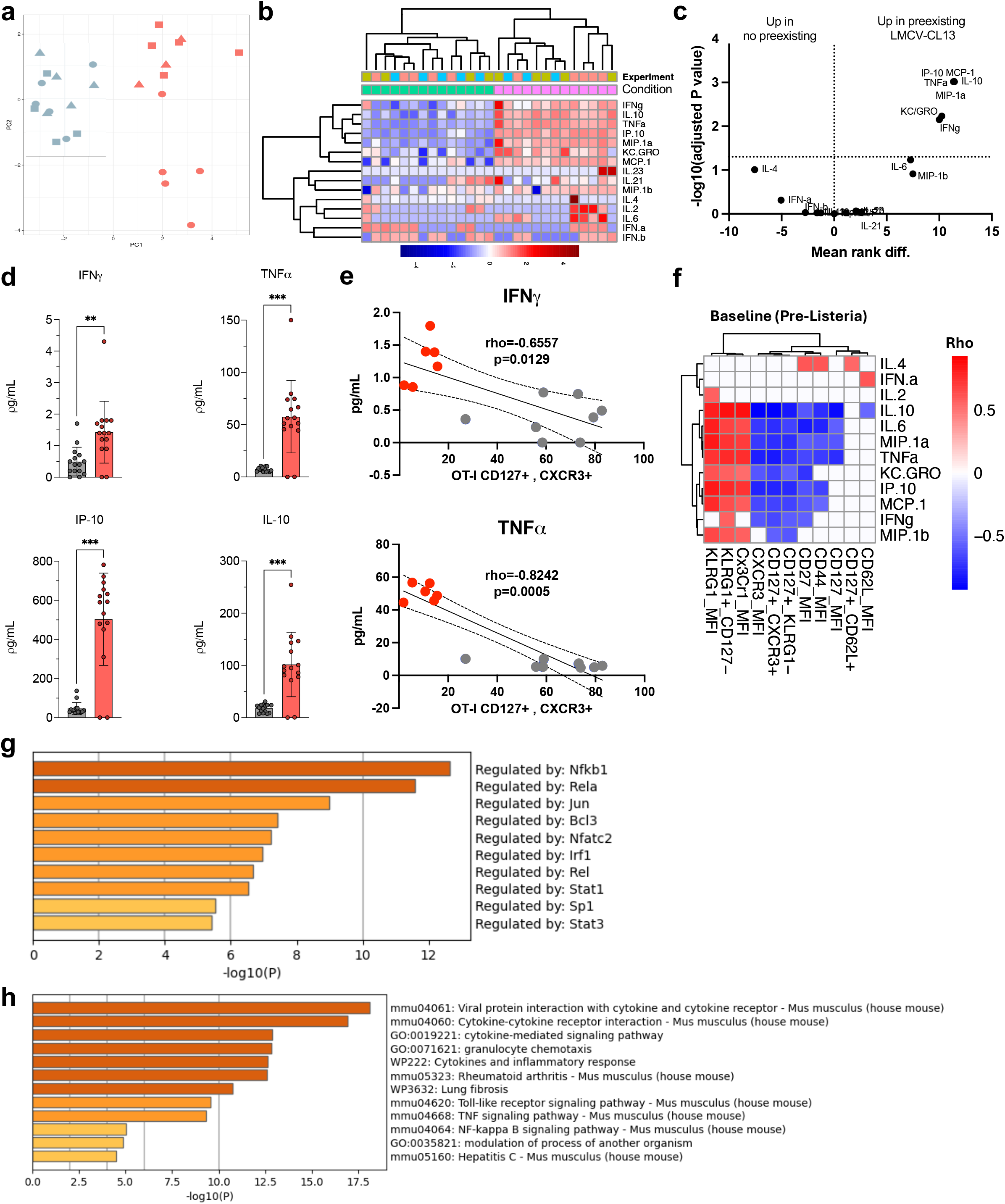
Preexisting persistent inflammation in chronically infected mice directly correlates with impaired memory CD8 T cell formation to subsequent acute infection. (A) Principal component analysis (PCA) of the levels of measured cytokines in the serum from mice with preexisting chronic LCMV-CL13 compared to control mice with no preexisting infection at baseline timepoint (1-3 days before *Listeria* infection). (B) Heatmap showing the levels of expression for all measured 15 cytokines at the baseline in mice from both groups. (C) Volcano plot showing significantly induced cytokines at baseline in mice with preexisting chronic LCMV-CL13 (right) compared to control mice with no preexisting infection (left). (Mann-Whitney U-test; all pass adj. p <0.05). (D) Serum level of expression of gamma-interferon (IFNg), tumor necrosis factor alpha (TNFa), IP-10, and interleukin 10 (IL-10) at baseline timepoint (1-3 days before *Listeria* infection). (E) Top, Spearman correlation of baseline IFNg levels and the percentage of antigen-specific CD8 T cells with a memory phenotype at d30. Bottom, Spearman correlation of baseline TNF levels and the percentage of antigen-specific CD8 T cells with a memory phenotype at d30. (F) Heatmaps of all significant correlations between flow cytometry features of antigen-specific OT-I CD8 T cells at d30 and baseline cytokine levels. (Spearman correlation; p <0.05) (G) Transcription factors predicted to drive the cytokine signature (listed above) using TRRUST; NFkB1/Rela/Rel are all driven by NF-kB; IRF1 and STAT1 driven by IFN; STAT3 driven by IL-6 and IL-10; and Jun induced by chemokines. (H) Top gene ontology (GO) pathways enriched for in the preexisting CL13 cytokine signature, including rheumatoid arthritis and lung fibrosis signatures.

Our analyses using TRRUST predicted enrichment of processes regulated by NFkB1, Rela, and Rel all driven by NF-κB; IRF1 and STAT1 driven by type I IFN; STAT3 driven by IL-6 and IL-10; and Jun induced by chemokines (Fig.4g). Gene ontology (GO) analyses, showed enrichment of numerous pathways in the preexisting CL13 cytokine signature, including cytokines and inflammatory response, rheumatoid arthritis, and lung fibrosis signatures (Fig.4h).

Collectively, these data suggest that the skewed cytokine milieu at baseline in mice with preexisting chronic infection with LCMV-CL13 has a major implication on the memory differentiation and phenotype of OT-I CD8 T cells on d30.

### Skewed inflm.-T_MEM_ phenotype is associated with skewed chromatin accessibility

Our results so far indicated that preexisting chronic LCMV-CL13 infection skewed the phenotype and subsets dynamics of memory T cells generated in response to subsequent acute infection with Listeria (hereafter termed inflm.-T_MEM_). This significant skewing persisted long-term post clearance of Listeria, suggesting that either the skewed transcriptional state reported previously^22, 24^ is continuously maintained by persistent inflammation, or that a skewed epigenetic gene regulation program is responsible for imprinting this cell state. Previous studies showed compromised recall and protective responses of adoptively transferred OT-I cells that were generated in mice with preexisting chronic disease, even though they were adoptively transferred to naïve mice before the rechallenge^22, 24^, suggesting that it is not merely a transcriptional state maintained under persistent pro-inflammatory conditions, but rather an epigenetic imprint that is not lost upon removal of all chronic conditions when adoptively transferring the cells into new naïve hosts.

To investigate the impact of preexisting chronic infection on the epigenetic landscape of the main subsets of T_MEM_, we used the same experimental design as Fig.2a, and we sort-purified the CD127+ CD62L+ T_CM_ and CD127+ CD62L– T_EM_ subsets of Listeria-specific OT-I CD8 T cells (Fig.5a) and performed Chromatin accessibility profiling using ATACseq on these sorted cells, then performed preliminary analyses of the data.

**Figure 5:**
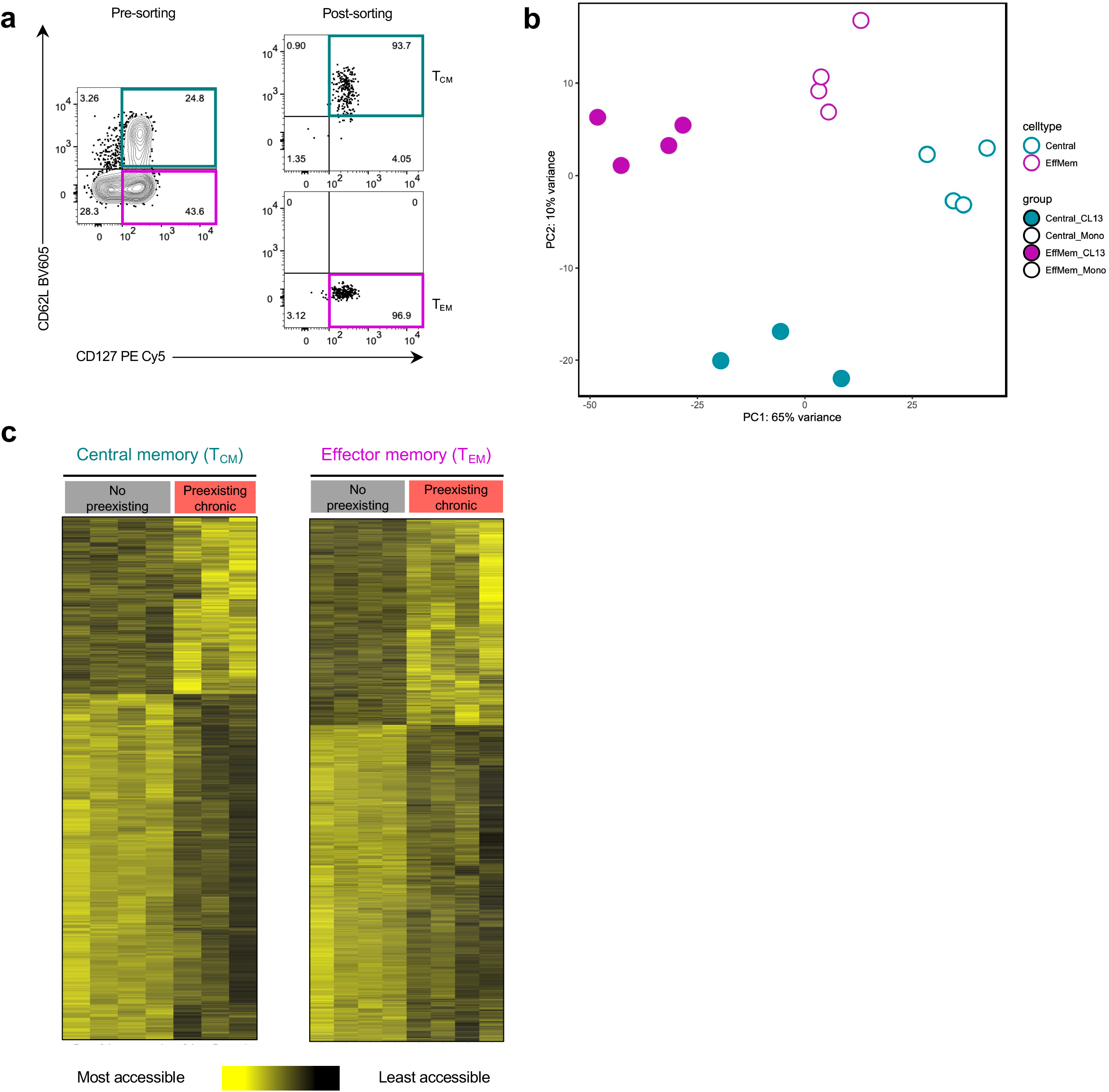
Preexisting chronic infection causes epigenetic skewing of both central memory (T_CM_) and effector memory (T_EM_) T cells. (A) Representative dot plots showing sorting purity of CD62L+ CD127+ central memory (T_CM_, top right) and CD62L– CD127+ effector memory (T_EM_, bottom right) phenotype, upon sorting LM-OVA specific OT-I CD8 T cells on d30 post Listeria infection (left). (B) Principal component analysis (PCA) of chromatin-accessibility profiles using ATAC-seq for OT-I T_CM_ and T_EM_ sort-purified at d30 post-Listeria infection (teal and purple colored, respectively), from mice with and without preexisting chronic LCMV-CL13 infection (solid and open circles, respectively). (C) Heatmaps of open chromatin regions (OCRs) significantly different between OT-I the group with and without preexisting chronic LCMV-CL13 infection, in central memory T cells (T_CM_, left) and effector memory T cells (T_CM_, left).

Principal component analysis (PCA) of our datasets showed a clear distinction between T_CM_ and T_EM_ isolated from the group with preexisting chronic infection compared to their counterparts from the control group (Fig.5b). The major principal component (PC1) that accounts for 65% of the differentially accessible chromatin regions (DACRs) mainly separated between the two conditions: hosts with preexisting chronic infection versus hosts with no preexisting infection.

Hundreds of genomic loci were differentially accessible chromatin regions (DACRs) between the two conditions (FDR<0.05) in the T_CM_ and T_EM_, respectively (Fig.5c). Notably, for both T_CM_ and T_EM_ from hosts with preexisting chronic infection, there was a higher number of DACRs that were less accessible compared to the control group, whereas lower number of DACRs that were more accessible.

Thus, our chromatin-accessibility landscape analyses of the two major subsets of memory T cells, central memory and effector memory, suggests that the compromised recall capacity observed by previous studies is due to major skewing of the epigenetic profile of both subsets.

### Skewed chromatin accessibility of memory T cells in hosts with preexisting chronic infection originated early during the effector phase

Previous studies reported that recall responses were compromised in OVA-specific OT-I cells from mice with preexisting chronic CLL, despite the fact that OT-Is were adoptively transferred into naïve mice just 7 days post mCMV-OVA infection and were left to recover in the naïve mice for 5 weeks before rechallenge^24^. This study also showed that the chromatin accessibility of total OT-Is from mice with preexisting chronic CLL were significantly different from OT-Is from naïve mice. However, this significant difference in the epigenetic profile is expected due to the significant difference in the composition of OT-I subsets between the two groups.

Whether epigenetic skewing originated early during the effector phase in the memory-precursor MPEC subset that mainly give rise to T_MEM_ is not known. To elucidate this, we sort-purified CD127+ KLRG1– OT-I cells on d8 post Listeria infection that represent the memory-precursor effector cells (MPECs) (Fig.6a), and performed Chromatin accessibility profiling using ATACseq on these sorted cells, then performed preliminary analyses of the data.

**Figure 6:**
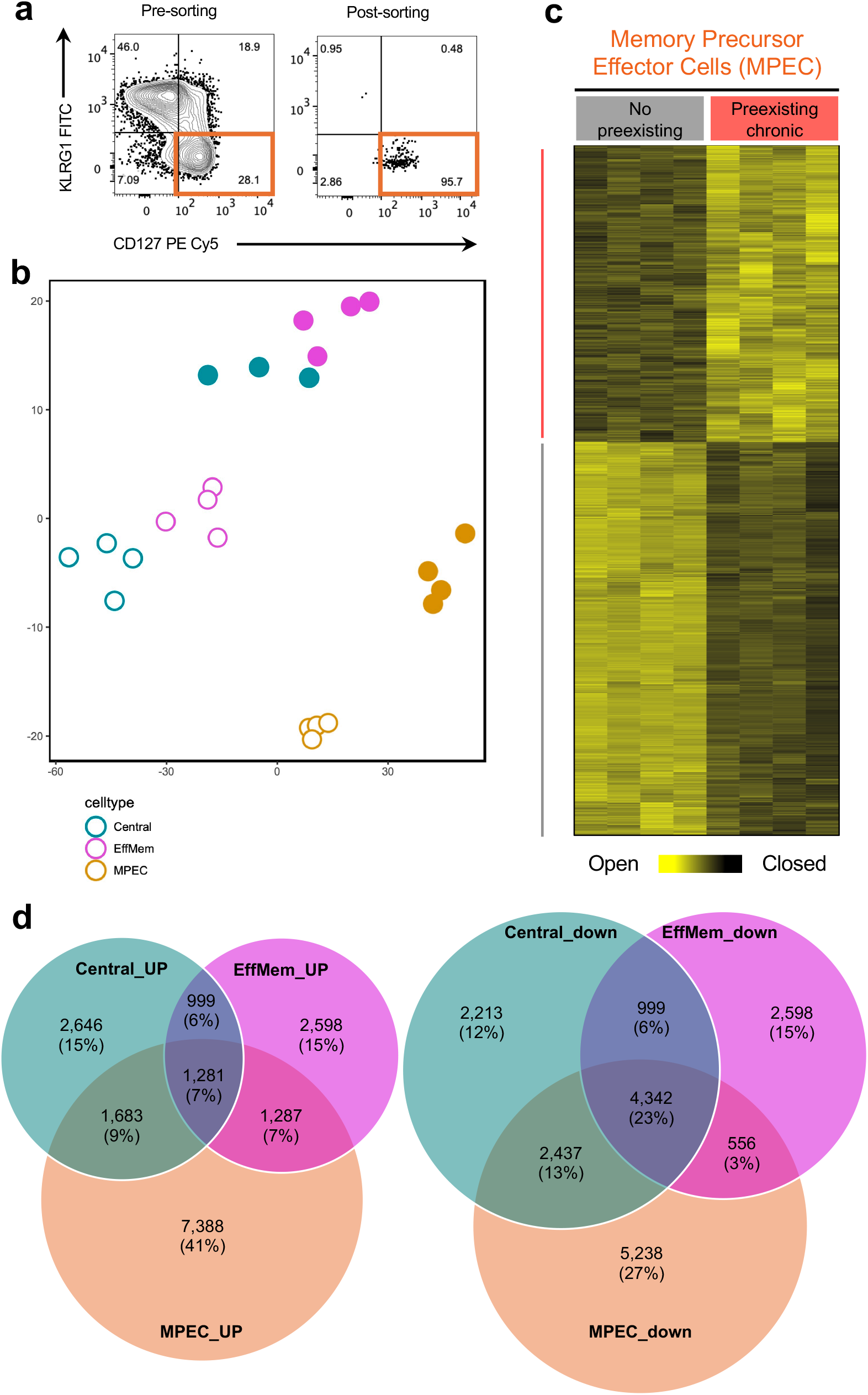
Epigenetic skewing is initiated early during the peak of the immune response in memory-precursor effector cells (MPECs). (A) Representative dot plots showing sorting purity of KLRG1– CD127+ memory precursor effector cells (MPEC, right) phenotype, upon sorting LM-OVA specific OT-I CD8 T cells on d8 post Listeria infection (left). (B) Principal component analysis (PCA) of chromatin-accessibility profiles using ATAC-seq for OT-I T_CM_ and T_EM_ sort-purified at d30 post-Listeria infection (teal and purple colored, respectively), and MPEC OT-I cells sort-purified at d8 post-Listeria infection (orange) from mice with and without preexisting chronic LCMV-CL13 infection (solid and open circles, respectively). (C) Heatmap of open chromatin regions (OCRs) significantly different between OT-I the group with and without preexisting chronic LCMV-CL13 infection, in KLRG1– CD127+ memory precursor effector T cells (MPEC) isolated and sort-purified at d8 post-Listeria infection. (D) Venn diagram of overlap between MPEC, T_CM_ and T_EM_ in OCRs significantly more accessible in the group with preexisting chronic LCMV-CL13 infection compared to the control group without preexisting chronic infection. (E) Venn diagram of overlap between MPEC, T_CM_ and T_EM_ in OCRs significantly less accessible in the group with preexisting chronic LCMV-CL13 infection compared to the control group without preexisting chronic infection.

Indeed, similar to the distinct profiles of T_CM_ and T_EM_ observed between the group with preexisting chronic infection and the control group (Fig.5b), the profiles of MPECs was distinct between the two groups (Fig.6b). Again, hundreds of genomic loci were differentially accessible chromatin regions (DACRs) between the two conditions (FDR<0.05) in the MPECs (Fig.6c). Importantly, there was a significant overlap in the DACRs that were significantly more accessible in T_CM_ and T_EM_ in the group with preexisting chronic infection and were already more accessible in MPECs, 16% and 14%, respectively. Even more significant was the overlap of DACRs that were significantly less accessible, where T_CM_ had 36% overlap with MPECS and T_EM_ had 26% overlap with MPECS.

Together, these data highlight that despite the lack of significant difference in effector functions at the peak of CD8 T cell responses to Listeria in mice with preexisting CL13, the skewing of the epigenetic landscape has already started during these early stages, and further accumulated during the differentiation of cells into the memory state.

## Discussion

We examined the differentiation of antigen-specific CD8 T cells generated to acute infection in hosts with preexisting chronic disease. During the early effector phase antigen-specific CD8 T cells (OT-I) generated to *Listeria monocytogenes* expressing ovalbumin (LM-OVA) demonstrated enrichment of short-lived effector cells (SLECs) in mice with preexisting chronic LCMV-CL13. However, overall cytokine production and polyfunctionality was not profoundly impacted, which resulted in similar clearance of Listeria. At later timepoints post LM clearance significant skewing of OT-I phenotype and T_CM_/T_EM_ dynamics was observed for T_MEM_ in mice with preexisting chronic infection (hereafter inflm.-T_MEM_), where the formation of T_CM_ was specifically compromised. Importantly, this was associated with a significantly altered chromatin-accessibility landscape for both T_CM_ and T_EM_, and this epigenetic skewing started early in the memory-precursor effector cells (MPECs). Differentially accessible chromatin regions (DACRs) that are more accessible in inflm.-T_MEM_ were enriched in genomic loci associated with transcription factors linked to proinflammatory cytokines, e.g. interferon regulatory factors (IRFs). This was suggested to be driven by a skewed cytokine milieu enriched in proinflammatory cytokines.

Two previous studies evaluated the “bystander” effect of chronic disease on CD8 T cell differentiation using mouse models^22, 24^. Martens et al. focused on the impact of preexisting chronic lymphocytic leukemia (CLL) on early time points at the peak of the effector OT-I response to mouse CMV expressing OVA (mCMV-OVA)^24^. Whereas Stelekati et al. focused on the impact preexisting chronic infection with LCMV-CL13, toxoplasma, or helminths on the transition of OT-I against VSV-OVA or LM-OVA from the effector to memory state^22^. For the latter study, in most experiments naïve OT-I were primed under normal conditions in naïve mice, then adoptively transferred into mice with preexisting chronic infection ^22^. Our results corroborate with the overall phenotypic conclusions from both studies and extends their findings, where we used the different gating strategies that would identify T_CM_/T_EM_, e.g. CD127/CD62L, or CD27/Cx3cr1. Another advantage is that we tracked the differentiation of OT-I within the same host since the initiation of infection, thus examining all imprints impacting the differentiation of OT-I throughout all phases of the subsequent acute infection. Additionally, this is more representative of the real-life situation, where the cells are primed and differentiate under the influence of the preexisting chronic disease conditions. Thus, our study design enabled capturing all imprints occurring since the earliest stages of priming, as well as imprints that keep accumulating throughout the various differentiation stages, while some of these imprints could have been missed by both studies, where the first study by Martens et al. only captured the early imprints, while Stelekati et al. only captured imprints that take place post the early priming and activation phase.

Of interest, OT-I in both studies showed compromised expansion when adoptively transferred into naïve mice and rechallenged with the antigen^22, 24^, even when T_EFF_ OT-I were transferred into naïve mice just 7 days post mCMV-OVA infection and were left to recover in the naïve mice for 5 weeks before rechallenging them. Stelekati et al. showed similar results, however, despite showing skewed phenotype and recall for these OT-I cells, recent epigenetic studies suggest that distinct epigenetic modules are imprinted in T cells during the very early stages of immune responses^27, 28, 29, 30^. Indeed, our ATACseq analyses demonstrating the skewed chromatin-accessibility profile of MPECs argue that studying the differentiation of antigen-specific T cells since the initial priming steps from the naïve to the effector state under conditions of preexisting chronic infection revealed many significant changes that became fixed in T_CM_ and/or T_EM_., with more epigenetic changes that became imprinted in T_CM_/T_EM_ as OT-I continued differentiating under preexisting chronic conditions. Recent studies suggest that preexisting infection impacts naïve “bystander” T lymphocyte repertoire even before encountering their cognate antigen^31^.

Epigenetic profiling showed great value inferring the history of cell differentiation through the imprints carved at different epigenomic loci, as well as predicting the future behavior of immune cells in various contexts^27, 32^. Our ATACseq analyses showed a stark difference in the chromatin-accessibility profile of Listeria-specific inflm-T_MEM_ from mice with preexisting chronic infection compared to T_MEM_ from mice with no preexisting infection (Fig. 5-6). Our chromatin-accessibility analysis identified the IRF family of transcription factors (TFs) as one of the top differential TFs in accessible chromatin at their binding sites, thus the transcription factor binding site (TFBS) analysis suggests that IRFs are involved in the skewed profile of inlfm-T_MEM_. Previous studies showed that knocking-out IRF2 in CD8 T cells in tumor settings resulted in uncoupling of the IFN-associated dysfunction within exhausted T cell (T_EX_) while preserving the cell priming and activation effects of IFN-I ^33^. This was associated with extensive reprogramming of T cell transcriptional networks and reshaping their chromatin accessibility landscape.

Chronic viral infections such as HIV, HBV, and HCV are associated with elevated levels of proinflammatory cytokines, such as TNF, IFN-I, and IL-6^34, 35, 36, 37^. LCMV-CL13 recapitulates the general proinflammatory profiles in these human chronic viral infections, where persistently elevated IFN-I was suggested as one of the drivers of the dysfunction of LCMV-specific T cells that could be reversed by blockade of IFN or IFN-receptor (IFNAR)-KO^38, 39^, and signatures of proinflammatory cytokines are enriched in exhausted T cells in LCMV-CL13 chronic setting^29, 32, 40^. Transcriptional profiling of CD8 inflm.-Tmem with preexisting chronic LCMV-CL13 showed enrichment of proinflammatory signatures, including IL-12, IFN-α, TNF and IL-6^22^. Modulating one of these inflammatory pathways, specifically knocking out type I interferon receptor alpha subunit (IFNAR-KO), partially restored cellular profile and recall capacity of T_MEM_ observed in mice with no preexisting infection^22^. Inflammatory mediators were shown to directly impact T_MEM_ phenotype, e.g. exposure of T cells to IL-6 or IFN-I caused downregulation of CD127 (the IL-7 receptor a subunit) and diminished IL-7 induced signaling and proliferation (reviewed in^34^). Additionally, T-bet, a TF downstream the proinflammatory JAK-STAT pathway that skews T cells towards SLECs on the expense of long-lived MPECs^23, 41^, was shown to be enriched in inflm.-T_MEM22_. Our data confirmed that the skewed cytokine milieu in the serum of mice with preexisting CL13 is directly correlated with the compromised memory differentiation of LM-OVA-specific OT-I CD8 T cells, and the reduced expression of memory markers on these cells.

Our study instigates several important translational and basic immunology questions. First, how can fine-tuning proinflammatory signaling in the context of preexisting infection therapeutically modulate inflm.-T_MEM_ differentiation? This necessitates understanding the mechanisms that modulates the chromatin-accessibility landscape. Recent data suggest that even in hosts with no preexisting chronic disease or persistent inflammation, early blockade of proinflammatory pathways would enrich the differentiation of T_MEM_ that would possess superior recall capacity^42^. Second, how does chronic disease impact innate immune cells, and consequently the priming of adaptive immune cells, and whether this is one of the underlying causes of epigenetic skewing of inflm.-T_MEM_? Third, would antigen-specific B cells and CD4 T cells be subject to similar skewing? Recent studies reported skewing of naïve CD4 T cells with preexisting helminths infection^31^, and memory B cells (B_MEM_) exhibited a cluster unique to chronic conditions driven by IFN-I^43^. Finally, our results for inflm.-T_MEM_ are in the context of preexisting systemic infection with LCMV-CL13, similarly the TCL1 model of CLL used by Martens et al., but it would be essential to understand how would localized solid tumors that are associated with heightened systemic inflammation impact responses to vaccines. Better understanding of mechanisms underlying skewed programming of inflm.-T_MEM_ in the different preexisting chronic contexts would enable novel therapeutic approaches to enhance vaccine-induced memory in these populations.

## Acknowledgments

This work was supported by an Emory MP3 Seed Grant (121861), and Department of Pathology and Laboratory Medicine Start-up funds to MSA. The authors would like to thank members of the Hakeem Lab, as well as members of the Pathology Advanced Translational Research Unit (PATRU) and Emory Vaccine Center (EVC) for their valuable discussions and comments.

We would also like to thank Dr. Mandy Ford and Ford Lab members for providing technical support with the Listeria monocytogenes and OT-I mouse colony set-up.

We would also like to thank the Emory Flow Cytometry Core (EFCC) and the Emory Integrated Genomics Core (EIGC) for their support. This study was supported in part by the Emory Flow Cytometry Core (EFCC), one of the Emory Integrated Core Facilities (EICF), and is subsidized by the Emory University School of Medicine. Additional support to EFCC was provided by the National Center for Georgia Clinical & Translational Science Alliance of the National Institutes of Health under Award Number UL1TR002378.

Research reported in this publication was supported in part by the Emory Integrated Genomics Core (EIGC) shared resource of Winship Cancer Institute of Emory University and NIH/NCI under award number P30CA138292.

We thank the NIH Tetramer Core Facility (contract number 75N93020D00005) for providing tetramers used in this study (H2-Kb OVA257-264, SIINFEKL). The NIH Tetramer Core Facility is supported through National Institute of Allergy and Infectious Diseases (NIAID) and co-funding from the National Cancer Institute (NCI). The content is solely the responsibility of the authors and does not necessarily represent the official views of the National Institutes of Health.

## Author contributions

MSA conceived and designed the study and experimental design. ASD performed the experiments, with help from EW, SJ and AO. ASD and EW were responsible for mouse breeding and colony maintenance with help from SJ. YMZ performed initial preparation and alignment for ATACseq data. Blaze Informatics performed bioinformatic analyses for ATACseq data. TG performed cytokine measurements. ASD performed flow cytometry and data analyses and prepared the figures. MSA wrote the manuscript. ASD wrote the M&M section. JAT supervised cytokine measurements and analyses, and contributed to manuscript proof-reading. SAR helped with data interpretation and manuscript proof-reading.

## Notes

### Competing Interest Statement

The authors have declared no competing interest.

